## Supplementary Figures for "Deep learning the collisional cross sections of the peptide universe from a million training samples"

**Supplementary Figures 1-5**

### Deep learning the collisional cross sections of the peptide universe from a million training samples

Florian Meier[1,#], Niklas D. Köhler[2,#], Andreas-David Brunner[1,#], Jean-Marc H. Wanka[2], Eugenia Voytik[1], Maximilian T. Strauss[1], Fabian J. Theis[2,3*] and Matthias Mann[1,4,*]

[1] Max Planck Institute of Biochemistry, Department Proteomics and Signal Transduction, Martinsried, Germany
[2] Helmholtz Zentrum München—German Research Center for Environmental Health, Institute of Computational Biology, Neuherberg, Germany
[3] Department of Mathematics, TU München, Munich, Germany
[4] NNF Center for Protein Research, Faculty of Health Sciences, University of Copenhagen, Copenhagen, Denmark

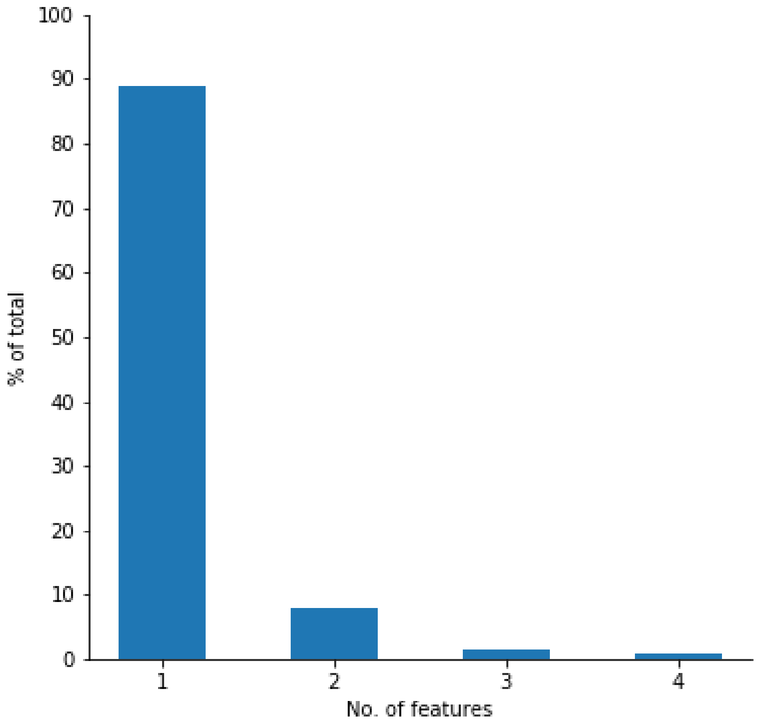


**Supplementary Figure 1**. Number of detected features per modified peptide sequence and charge state in single LC-TIMS-MS experiments.

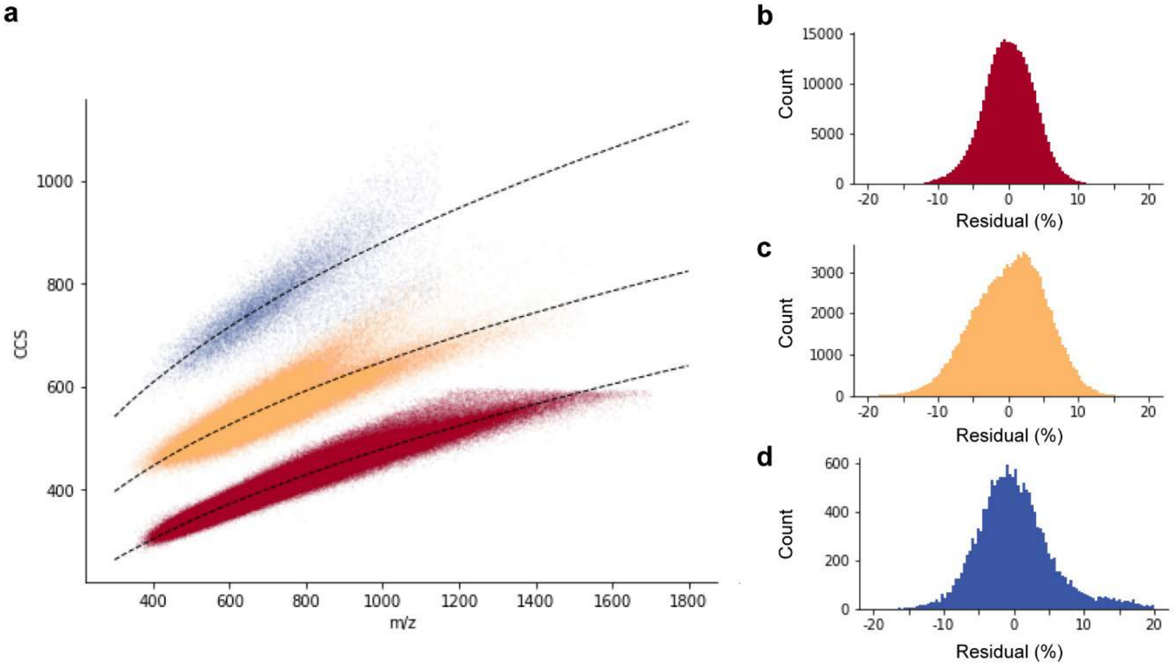


**Supplementary Figure 2**. **a,** Distribution of tryptic peptides in the *m/z* vs. CCS space color-coded by charge state as in Figure 1. Fitted power-law (A*x^b)) trend lines (dashed lines) visualize the correlation of ion mass and mobility in each charge state. **b-d,** Residuals for charge states 2, 3 and 4.

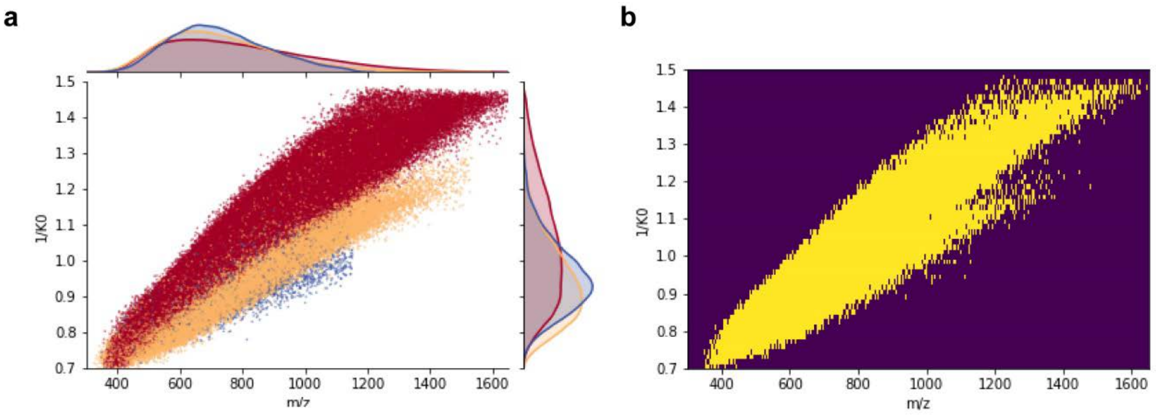


**Supplementary Figure 3**. **a,** Distribution of tryptic peptides in the *m/z* vs. ion mobility ($1/K_0$) space color-coded by charge state as in Figure 1. **b,** Estimating the peak capacity (Φ) of two-dimensional peptide separation with TIMS-MS. In an ideally orthogonal 2D separation, the total peak capacity would be $\Phi_{MS} * \Phi_{TIMS}$. Assuming an ion mobility resolution of 60 ($(1/K_0) / \Delta(1/K_0)$), the average peak full width at half maximum is 0.018 Vs $cm^{-2}$ in the peptide $1/K_0$ range (0.7-1.5 Vs $cm^{-2}$). This would result in a theoretical peak capacity of $\Phi_{MS}$ * 44. However, the correlation of mass and mobility reduces the effective peak capacity and the 2D histogram analysis (1350 *m/z* x 44 ion mobility bins) shows that 96% of the peptides occupy about 27% of the total area (yellow vs. purple area). Using this as a correction factor, we estimate the peak capacity of TIMS-MS to about $\Phi_{MS}$ * 12.

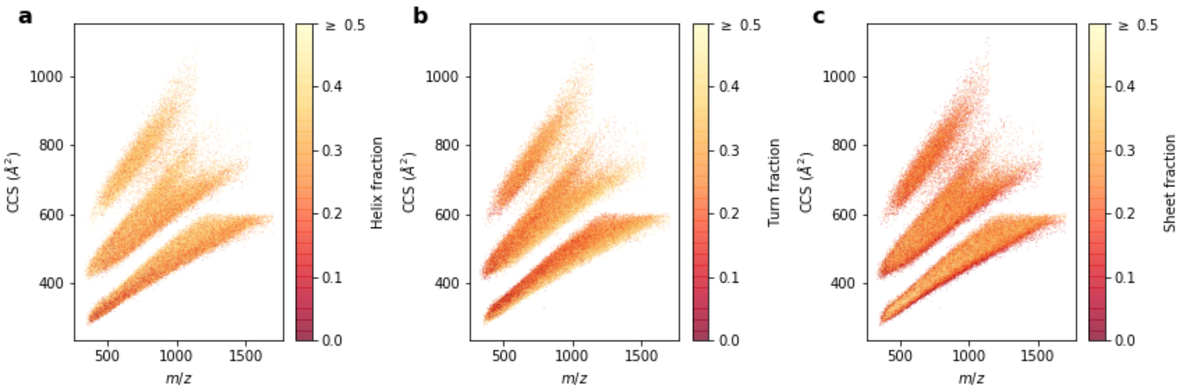


**Supplementary Figure 4**. Fraction of amino acids favoring **a**, helical (A, L, M, H, Q, E), **b**, turn (V, I, F, T, Y) and **c**, sheet (G, S, D, N, P) secondary peptide structures according to ref.[42].

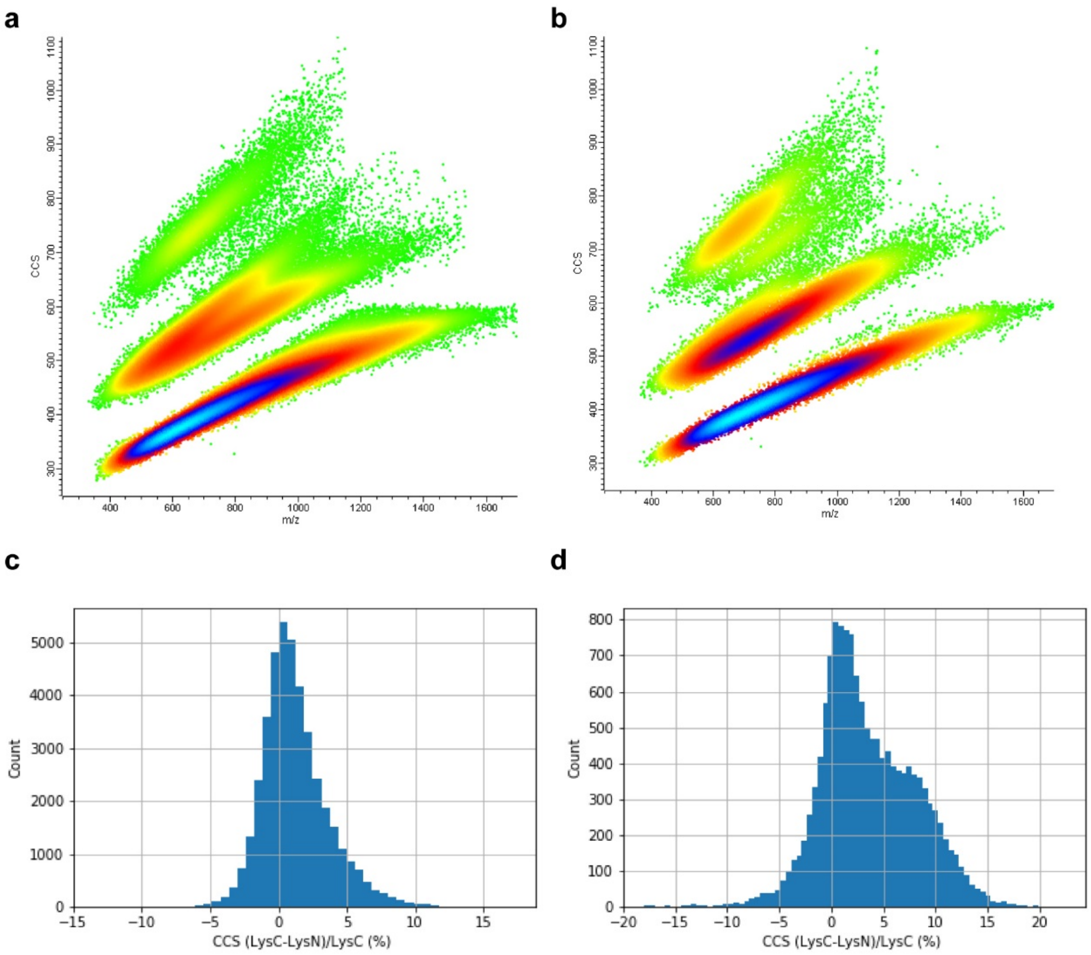


**Supplementary Figure 5.** CCS comparison of LysC and LysN digests. **a**, density distribution of LysC peptides. **b**, Density distribution of LysN peptides. **c**, Pairwise comparison of doubly-charged peptides with the same internal sequence. **d**, Pairwise comparison of triply-charged peptides with the same internal sequence.
